# Time-Resolved Profiling: Metabolic Adaptation & Morphology-Specific Drug Response in MCF-7 spheroids

**DOI:** 10.64898/2026.07.24.740505

**Authors:** Yaping Li, Nora Schäfer, Sylwia Barker, Marcel Utz, Annamarija Raic

## Abstract

Three-dimensional (3D) tumor models exhibit drug responses that differ from conventional 2D cultures. However, how cellular metabolism dynamically evolves across culture models and during drug treatment remains poorly understood. We compared the responses of MCF-7 cells in 2D and 3D environments to 5-fluorouracil (5-FU), combining ¹H NMR cellular metabolomics with viable cell counts, the GLUT1-positive population, gene expression, ATP activity, and morphology. Principal component analysis revealed that culture dimensionality, rather than 5-FU treatment, was the primary driver of metabolic flux variation. 3D spheroids exhibited higher glycolytic flux at 72h. Importantly, this elevated glycolysis reflected a higher per-cell flux in larger spheroids rather than an increased cell number. We further observed a higher proportion of GLUT1-positive cells and increased HK2 expression in 3D culture, together with an epithelial phenotype characterized by increased CDH1 and decreased VIM expression. Functionally, 3D displayed maintaining higher cell viability, ATP activity following treatment. Together, these findings suggest that 3D architecture promotes a metabolically defensive phenotype and cellular metabolic behaviors is associated with morphology, which may inform future drug screening model selection.

**Blurb:** Time-resolved NMR metabolomics reveals that 3D culture architecture, rather than 5-FU treatment, defines the metabolic phenotype of MCF-7 cells, linking spheroid morphology with per-cell glycolytic activity, ATP preservation, and reduced chemotherapy sensitivity.

**Synopsis:** 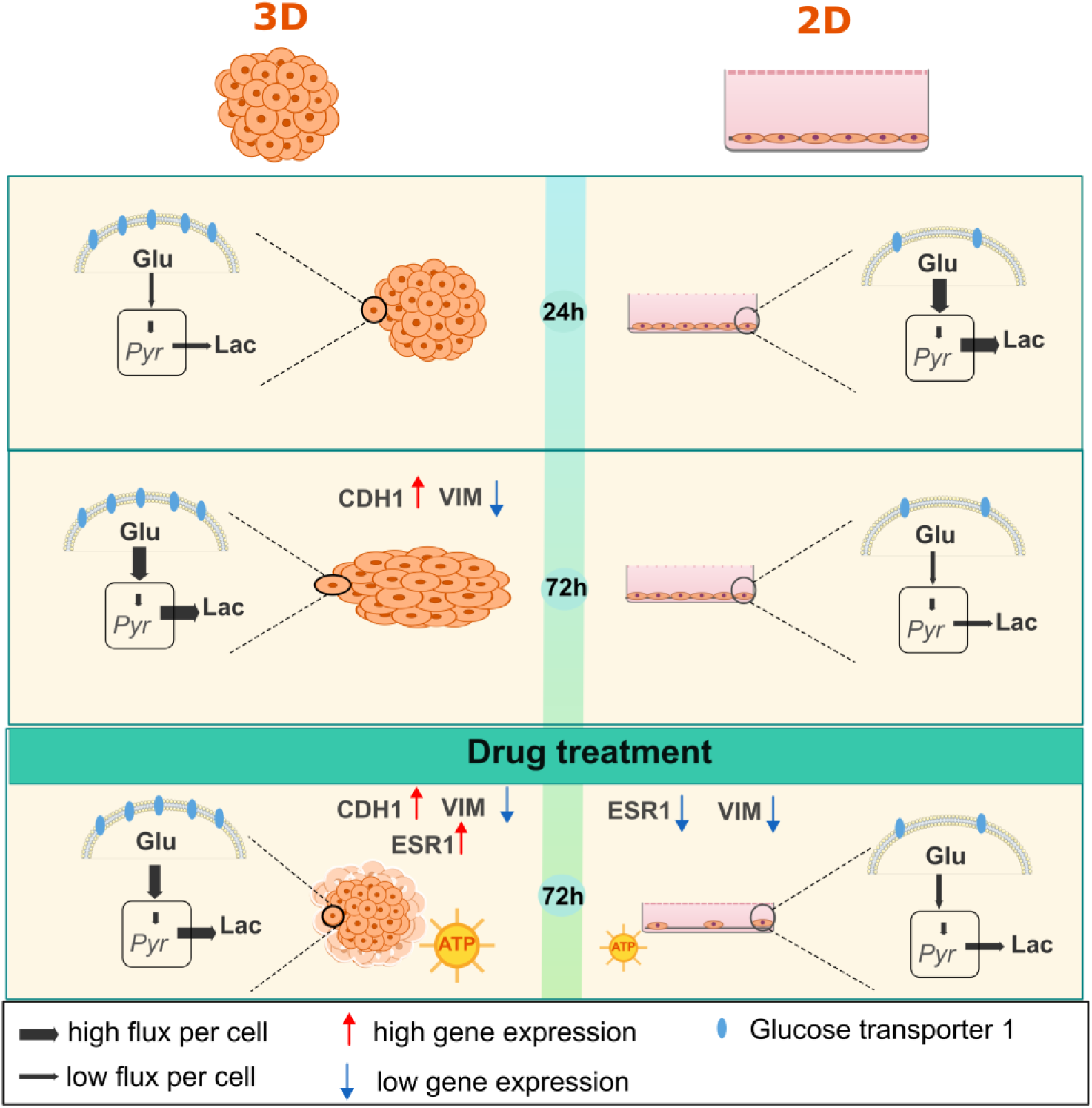

Graphical abstract — Time-resolved metabolic and phenotypic responses of 2D and 3D MCF-7 cultures to 5-fluorouracil.

**Bullet points:**

- Time-resolved ¹H NMR cellular metabolomics combined with multilevel phenotypic readouts reveals that culture dimensionality, rather than 5-FU chemotherapy, is the dominant determinant of metabolic phenotype in MCF-7 breast cancer cells.
- 3D spheroids develop a glycolysis-dominant metabolic state with markedly elevated GLUT1⁺ populations and HK2 transcript, accompanied by substantially reduced sensitivity to 5-FU compared with 2D monolayers.
- Spheroid morphology correlates with cellular metabolic flux: larger and more elongated spheroids exhibit higher glycolytic activity on a per cell, independent of spheroid cell number.
- The 3D-defined metabolic state buffers ATP under 5-FU stress and reinforces an epithelial transcriptional program (CDH1↑, VIM↓), arguing that culture architecture must be considered when interpreting preclinical drug responses.

## 1. Introduction

This study investigated how 2D and 3D breast cancer cultures differ in their metabolic response to 5-fluorouracil (5-FU), and how these metabolic changes relate to cellular phenotype. Breast cancer remains the leading malignancy among women (Bray *et al*, 2024), and the in vitro models used in preclinical drug screening directly shape which therapeutic candidates advance to patients. However, the gap between 2D monolayer responses and in vivo outcomes has become a recognized bottleneck (Hammond *et al*, 2024). Three-dimensional spheroid models have therefore been increasingly adopted to recapitulate the in vivo tumor microenvironment (Zanoni *et al*, 2016; Dhandapani *et al*, 2023), and ^1^H NMR-based cellular metabolomics has emerged as a label-free route to profile their metabolic state (Beckonert *et al*, 2007; Patra *et al*, 2021). Despite these advances, how culture architecture jointly reshapes metabolic flux, cellular phenotype, and drug responsiveness over time remains insufficiently understood, in part because most studies report without integrating metabolism with phenotypic readouts. To address this gap, we quantified metabolic flux in 2D and 3D MCF-7 cultures using ¹H NMR metabolomics and integrated these data with multilevel phenotypic characterization over 72 h of 5-FU treatment.

Breast cancer remains the most common malignancy among women worldwide, with approximately 2.3 million new cases diagnosed annually and accounting for nearly 15% of female cancer-related deaths (Bray *et al*, 2024). Despite extensive efforts in preclinical drug development, conventional animal models often fail to accurately predict human therapeutic responses due to interspecies differences (Loewa *et al*, 2023). Consequently, in vitro culture systems remain essential tools for cancer research and drug screening. Traditional two-dimensional (2D) monolayer cultures are widely used due to their simplicity and reproducibility; however, they fail to recapitulate key features of the in vivo tumor microenvironment, including cell–cell and cell– extracellular matrix (ECM) interactions (Saraswathibhatla *et al*, 2023), polarity, and nutrient and oxygen gradients (Edmondson *et al*, 2014). In contrast, three-dimensional (3D) spheroid models better recapitulate the architecture and microenvironment of solid tumors, providing a more physiologically relevant platform for studying tumor biology (Zanoni *et al*, 2016; Nunes *et al*, 2019; Dhandapani *et al*, 2023). Importantly, 3D culture has been shown to profoundly influence tumor phenotypes, including proliferation (Dhandapani *et al*, 2023), invasion (Chong *et al*, 2024), stemness (Ghanbari Movahed *et al*, 2023), and drug responsiveness (Crispim *et al*, 2025; Kes *et al*, 2024).

One hallmark of 3D spheroids is the extensive cell–cell contacts that promote adherens junction formation, intercellular signaling, and the maintenance of epithelial characteristics that are difficult to reproduce in monolayer cultures (Mangani *et al*, 2025). In breast cancer models, spheroid cultures are consistently associated with reduced sensitivity to cytotoxic agents compared with 2D monolayers (Breslin & O’Driscoll, 2016), among which enhanced cell–cell interactions and integrin-mediated signaling represent important contributing factors (Lovitt *et al*, 2018). 3D organization has been shown to preserve subtype-specific tumor features that are largely absent in 2D cultures, as demonstrated in MCF-7 spheroid models (Dhandapani *et al*, 2023). However, despite growing understanding of the signaling and phenotypic consequences of enhanced cell– cell interactions in 3D systems, the metabolic mechanisms underlying 3D culture–dependent drug responses remain poorly understood.

Metabolic reprogramming is a hallmark of cancer and plays a central role in tumor progression and therapeutic resistance (Pavlova *et al*, 2022). Cancer cells frequently exhibit aerobic glycolysis (the Warburg effect) together with rewired energy and nutrient metabolism to support biosynthetic demands (Liberti & Locasale, 2016; Faubert *et al*, 2020). Importantly, metabolic plasticity enables tumor cells to adapt to therapeutic stress and is closely linked to drug resistance (Tidwell *et al*, 2022; Wang *et al*, 2021). However, despite growing evidence that 3D culture systems exhibit distinct metabolic features and altered drug responsiveness, the metabolic mechanisms linking 3D architecture to drug response remain insufficiently understood. Conventional mass spectrometry– based metabolomics requires cell lysis and metabolite extraction, thereby disrupting 3D spatial organization and preventing longitudinal metabolic monitoring. In contrast, proton nuclear magnetic resonance (^1^H NMR) spectroscopy enables label-free and quantitative analysis of cellular metabolites in spent media and has recently been applied to 3D spheroid systems (Beckonert *et al*, 2007; Patra *et al*, 2021; Palma *et al*, 2016). Despite these advances, how 3D architecture shapes dynamic metabolic flux over time and how these metabolic adaptations relate to cellular phenotype and drug responsiveness remain poorly understood.

Here, using breast cancer cells as a model, we investigate how culture architecture influences metabolic flux and cellular responses to 5-fluorouracil (5-FU). We combine time-resolved ¹H NMR-based metabolic flux profiling with multilevel phenotypic characterization to capture metabolic and phenotypic changes across 2D and 3D cultures. The results reveal that culture architecture is the major driver of metabolic variation, whereas drug-induced metabolic alterations depend on the culture context. Single-cell metabolic activity within spheroids is associated with spheroid morphology, accompanied by culture-dependent differences in proliferation and GLUT1 expression. Together, these findings highlight the importance of considering culture architecture in metabolic studies and provide a basis for developing more physiologically relevant models for drug evaluation.

## 2. Methods

## Reagents and Tools Table

| Reagent/Resource | Reference or Source | Identifier or Catalog Number |
| --- | --- | --- |
| Experimental models |  |  |
| MCF-7 human breast cancer cells | Merck/Sigma-Aldrich | ECACC 86012803 |
| **Chemicals, peptides, and recombinant proteins** |  |  |
| DMEM, 4.5 g/L glucose, L-glutamine, sodium pyruvate | Corning | 10-013-CV |
| Fetal bovine serum (FBS) | Gibco | A5670701 |
| Penicillin–streptomycin | Gibco | 15140-122 |
| Trypsin–EDTA (0.25%) | Gibco | 25200-056 |
| Accutase | Bio&SELL | BS.2193 |
| 5-Fluorouracil (5-FU) | Sigma-Aldrich | F6627 |
| 3-(Trimethylsilyl) propionic-2,2,3,3-d<sub>4</sub> acid, sodium salt (TSP-d<sub>4</sub>) | Thermo Fisher Scientific | A14489.06 |
| Deuterium oxide (D<sub>2</sub>O) | Thermo Fisher Scientific | 166301000 |
| **Antibodies** |  |  |
| Anti-GLUT1, FITC-conjugated (clone GLUT1/2476) | Thermo Fisher Scientific | 18246776 |
| Mouse IgG2b κ isotype control, FITC | Thermo Fisher Scientific | 18264025 |
| **Commercial assays and kits** |  |  |
| CellTiter-Glo® 3D Cell Viability Assay | Promega | G9682 |
| LIVE/DEAD® Cell Imaging Kit (488/570) | Invitrogen | R37601 |
| LIVE/DEAD® Fixable Far-Red Dead Cell Stain | Invitrogen | L10120 |
| RNeasy Micro Kit | Qiagen | 74004 |
| iScript cDNA Synthesis Kit | Bio-Rad | 1708891 |
| iTaq Universal SYBR Green Supermix | Bio-Rad | 1725121 |
| PrimePCR SYBR Green assays (10 genes) | Bio-Rad | Assay IDs listed in Section 2.9 |
| **Consumables and other materials** |  |  |
| BIOFLOAT™ 96-well U-bottom ultra-low-attachment plate | faCellitate | F202003 |
| 5 mm NMR tubes | Wilmad | 6287509 |
| **Software and algorithms** |  |  |
| ImageJ/FIJI | NIH | RRID:SCR_003070 / SCR_002285 |
| Chenomx NMR Suite | Chenomx | RRID:SCR_015708 |
| Julia (v1.11 and v1.12) | Julia Language | RRID:SCR_021666 |
| R (v4.3.3) | R Foundation | RRID:SCR_001905 |
| CFX Maestro Software | Bio-Rad | RRID:SCR_018900 |
| Floreada flow cytometry analysis platform | Floreada | https://floreada.io/ |
| **Instruments** |  |  |
| JEOL 14.1 T JMTC-600/54/XC/WS NMR spectrometer (600 MHz) | JEOL |  |
| CFX Opus 96 Real-Time PCR System | Bio-Rad |  |
| MACSQuant flow cytometer | Miltenyi Biotec |  |
| SpectraMax i3x microplate reader | Molecular Devices |  |
| Olympus CKX53 inverted microscope | Olympus |  |
| EVE Plus Automated Cell Counter | NanoEntek |  |

### 2.1 Cell culture and spheroid formation

MCF-7 human luminal epithelial breast cancer cells were obtained from Merck/Sigma-Aldrich. Cells were maintained in Dulbecco’s Modified Eagle Medium (Corning) containing 4.5 g/L (25 mM) glucose, 4 mM L-glutamine, and 1 mM sodium pyruvate, supplemented with 10% fetal bovine serum (FBS; Gibco) and 1% (v/v) penicillin–streptomycin (Gibco), at 37 °C in a humidified atmosphere containing 5% CO₂. Cells were subcultured upon reaching approximately 80% confluency by enzymatic dissociation with 0.25% trypsin–EDTA solution (Gibco). For 2D monolayer cultures used in the treatment experiments, cells were seeded in 24-well plates at a density of 5 × 10⁵ cells/mL and allowed to attach overnight. For three-dimensional (3D) spheroid cultures, trypsinized single cells were resuspended in complete culture medium at 5 × 10⁵ cells/mL, and 100 µL of cell suspension was seeded per well into round-bottom ultra-low-attachment 96-well plates (BIOFLOAT™; faCellitate). Spheroids were allowed to form by self-aggregation overnight under standard culture conditions prior to treatment.

### 2.2 IC₅₀ determination

The half-maximal inhibitory concentration (IC₅₀) of 5-FU (Sigma-Aldrich) was determined using 2D monolayer cultures. Cells were seeded in 96-well plates at a density of 5 × 10⁵ cells/mL and allowed to attach overnight. The medium was replaced with DMEM containing 25 mM HEPES buffer and 5-FU at concentrations of 0, 100, 200, 400, 800, and 1600 µM, followed by incubation for 72 h at 37 °C in a humidified 5% CO₂ atmosphere. Cell viability was assessed using the CellTiter-Glo® 3D Cell Viability Assay (Promega). Briefly, 100 µL of CellTiter-Glo® reagent was added to each well, plates were mixed by pipetting for 5 min to induce cell lysis, and incubated for 20 min at room temperature in the dark to stabilize luminescence. Luminescence was measured using a SpectraMax i3x microplate reader (Molecular Devices). Three independent experiments were performed. Viability was expressed as a percentage relative to solvent controls, and IC₅₀ values were determined by nonlinear regression in Julia v1.11 (Supplementary Fig. S1).

### 2.3 5-FU treatment and sample collection

Based on the IC₅₀ results, a working concentration of 220 µM 5-FU (approximately the IC₅₀ value) was selected for all subsequent experiments. Monolayer and spheroid cultures were prepared as described in Section 2.1.For drug treatment, The 5-FU stock solution was added to DMEM supplemented with 25 mM HEPES to achieve the desired final concentrations. Untreated controls received the same HEPES-buffered DMEM without 5-FU. Samples were collected at 24, 48, and 72 h post-treatment. At each timepoint, culture supernatants were collected, centrifuged at 500 × g for 5 min to remove cell debris, transferred to 1.5 mL reaction tubes, and stored at −20 °C until NMR analysis. At each timepoint, cell-free blank controls (HEPES-buffered DMEM incubated under identical conditions without cells) were collected in parallel to serve as the baseline reference for NMR metabolite quantification, thereby accounting for any evaporation-induced concentration changes over time. Three independent biological replicates were performed.

### 2.4 Morphological analysis of spheroids

Brightfield images of spheroids were acquired at 0, 24, 48, and 72 h post-treatment using an Olympus CKX53 inverted microscope (Olympus) equipped with a 4× objective lens. Images were saved as JPEG files and analyzed using ImageJ/FIJI software (NIH). Morphological parameters including projected area, aspect ratio, and maximum Feret diameter (the longest distance between any two points on the object boundary) were quantified for each spheroid at each timepoint. A minimum of 12 spheroids per condition per timepoint were analyzed across three independent experiments.

### 2.5 Cell viability assessment

#### 2.5.1 LIVE/DEAD fluorescence imaging

Cell viability was visualized at 0 h and 72 h post-treatment using the LIVE/DEAD® Cell Imaging Kit (Invitrogen), which labels live cells with calcein AM (green fluorescence) and dead cells with BOBO-3 iodide (red fluorescence). Monolayer cultures and spheroids were incubated with 50 µL of staining solution in 50 µL PBS for 30 min at room temperature, washed once with PBS, and imaged using an inverted fluorescence microscope (Olympus CKX53). Green fluorescent area and mean fluorescence intensity were quantified using ImageJ/FIJI software to assess live cell retention and treatment response.

#### 2.5.2 ATP activity assay

Cellular ATP levels were quantified at 72 h post-treatment using the CellTiter-Glo® 3D Cell Viability Assay (Promega) according to the manufacturer’s instructions. This assay measures intracellular ATP as an indicator of metabolically active cells based on a luciferase-mediated bioluminescent reaction. Briefly, 100 µL of reagent was added to each well containing 100 µL of culture medium, plates were mixed by pipetting for 5 min to induce cell lysis, and incubated for 20 min at room temperature in the dark to stabilize luminescence. Luminescence was measured using a SpectraMax i3x microplate reader. Results were expressed as the ratio of 5-FU-treated to control luminescence for each culture type.

### 2.6 Flow cytometry

#### 2.6.1 Sample preparation and spheroid dissociation

At 24, 48, and 72 h post-treatment, monolayer cells were harvested by trypsinization (0.25% trypsin–EDTA). For spheroid cultures, gentle enzymatic dissociation was performed using Accutase (Bio&SELL) for 5 min at room temperature to obtain single-cell suspensions while preserving cell surface epitopes required for downstream GLUT1 staining. Single-cell suspensions from both culture systems were washed once with PBS and pelleted by centrifugation at 400 × g for 4 min.

#### 2.6.2 Cell viability and GLUT1 staining

Both cell viability and total GLUT1 protein abundance were assessed simultaneously from the same flow cytometric acquisition using a sequential staining protocol. Single-cell suspensions were first aliquoted (2 × 10⁵ cells per tube) and then incubated with LIVE/DEAD® Fixable Far-Red Dead Cell Stain (1 µg/mL; Invitrogen) in PBS for 30 min at room temperature to label dead cells. In live cells, this amine-reactive dye labels only cell-surface amines (dim fluorescence), whereas in dead cells it penetrates compromised membranes and additionally reacts with abundant intracellular amines (bright fluorescence), enabling discrimination between live and dead populations; the resulting covalent bonds ensure signal retention through subsequent fixation and permeabilization steps. After washing with PBS, cells were fixed with 3.7% paraformaldehyde (PFA) for 10 min at room temperature and washed once. Fixed cells were incubated in permeabilization buffer (PBS containing 0.1% triton-x100) for 15 min at room temperature, followed by incubation with a directly conjugated anti-GLUT1 antibody (clone GLUT1/2476, FITC-conjugated; Thermo Fisher Scientific) at a dilution of 1:50 for 1 h at room temperature. A matched isotype control (mouse IgG2b kappa, FITC; Thermo Fisher Scientific) was used at the same concentration to define the gating threshold for GLUT1-positive cells. After staining, cells were washed with PBS and analyzed immediately. All flow cytometry data were acquired on a MACSQuant flow cytometer (Miltenyi Biotec) and analyzed using the Floreada online platform (https://floreada.io/). A minimum of 10000 events were recorded per sample. Viable cell numbers were calculated by multiplying the total cell count obtained from the EVE Plus Automated Cell Counter (NanoEntek) by the live cell percentage determined from flow cytometric live/dead staining at each timepoint.

### 2.7 ¹H NMR spectroscopy and metabolite quantification

#### 2.7.1 Sample preparation

Culture media stored at −20 °C were thawed at room temperature prior to NMR analysis. NMR samples were prepared by mixing culture supernatant with D₂O (Thermo Fisher Scientific) containing 5 mM 3-(trimethylsilyl) propionic-2,2,3,3-d₄ acid sodium salt (TSP-d₄; Thermo Fisher Scientific), yielding a final TSP concentration of 0.5 mM. D₂O was used for field-frequency locking. Samples were transferred to 5 mm NMR tubes (Wilmad). Time-matched cell-free blank controls (Section 2.3) were prepared identically to establish baseline metabolite concentrations at each timepoint.

#### 2.7.2 Data acquisition

All ^1^H NMR measurements were carried out on a JEOL 14.1 T JMTC-600/54/XC/WS spectrometer (600 MHz for ^1^H). The 90° proton pulse was calibrated to 7.41 µs at 8.9 dB, corresponding to a nutation frequency of 33.7 kHz. Water was suppressed using a presaturation pulse sequence with the presaturation pulse power of 67 dB, corresponding to a nutation frequency of 42 Hz. 35,874 data points were acquired with 64 scans and 4 prescans with a relaxation delay of 5 s.

#### 2.7.3 Spectral processing and metabolite quantification

The NMR data were processed in Julia 1.12 with an NMR package developed by Marcel Utz (Utz, 2021). For spectral processing, the free induction decay (FID) signals were Fourier transformed and zero-filled to 2^17^ (131,072) data points, and an apodization function with 0.5 Hz line broadening was applied. Zero-order phase correction and baseline correction were performed. Metabolite identification was performed by comparison with the Human Metabolome Database (HMDB) and Chenomx NMR Suite spectral library; A total of 13 metabolites were identified and quantified: glucose, lactate, pyruvate, glutamine, glutamate, alanine, glycine, branched-chain amino acids (BCAAs: leucine, isoleucine, and valine), phenylalanine, tyrosine, histidine, formate, and acetate. Metabolite concentrations (mM) were determined by integration of characteristic resonance peaks (integration regions listed in Supplementary Table S1) relative to the TSP internal standard at 0 ppm (0.5 mM, 9 equivalent protons) using the following equation:

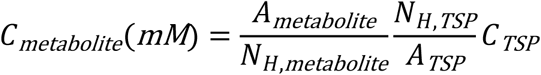

where A_metabolite denotes the integrated peak area, N_H,metabolite the number of equivalent protons contributing to the resonance, and C_TSP the known TSP concentration (0.5 mM, N_H,TSP = 9). A_TSP is the integral of the TSP peak.

### 2.8 Metabolic rate calculation

Average metabolic rates from seeding (t = 0 h) to each sampling timepoint (24, 48, 72 h) were calculated as:

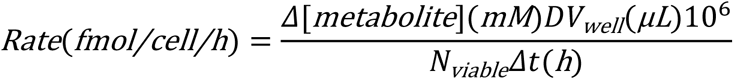

where Δ[metabolite] is the supernatant concentration change relative to a time-matched cell-free blank; D and V_well are the dilution factor and per-well culture volume; and N_viable is the per-well linear-integrated viable cell count (Matsuda *et al*, 2017):

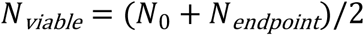

Both *N*_0_ and *N_endpoint_* refer to actual per-well counts, ensuring consistent units between numerator and denominator. Negative rates indicate net consumption; positive rates, net production. The lactate-to-glucose ratio was calculated as the lactate production rate divided by the absolute glucose consumption rate per condition × culture × timepoint × replicate; the theoretical maximum is 2.0 for complete glycolytic conversion of glucose to lactate.

### 2.9 RNA isolation and quantitative RT-PCR

Total RNA was extracted at 72 h post-treatment from both monolayer and spheroid cultures using the RNeasy Micro Kit (Qiagen) according to the manufacturer’s instructions. RNA concentration and purity were assessed by spectrophotometry (A260/A280 ratio ≥ 1.8). RNA samples were reverse-transcribed into complementary DNA (cDNA) using the iScript cDNA Synthesis Kit (Bio-Rad). Quantitative real-time PCR (qPCR) was performed using the CFX Opus 96 Real-Time PCR System (Bio-Rad) with iTaq Universal SYBR Green Supermix (Bio-Rad) and gene-specific PrimePCR assays (Bio-Rad), consisting of unlabeled primer pairs. Thermocycling conditions were as follows: initial activation at 95 °C for 2 min, followed by 39 cycles of denaturation at 95 °C for 5 s and annealing/extension at 60 °C for 30 s. A melt curve analysis was performed from 65 °C to 95 °C in 0.5 °C increments with 5 s per step to confirm amplification specificity. Nine target genes were analyzed spanning glycolytic enzymes (HK2, hexokinase 2; PKM2, pyruvate kinase M2), glutaminolysis (GLS1, glutaminase 1), hypoxia response (HIF-1α, hypoxia-inducible factor 1α), epithelial–mesenchymal transition (EMT) markers (CDH1, E-cadherin; VIM, vimentin), hormone receptor and differentiation markers (ESR1, estrogen receptor α; GATA3, GATA binding protein 3), and tumor suppression (TP53, tumor protein p53). The following PrimePCR assays (Bio-Rad) were used: GAPDH (Assay ID: qHsaCED0038674), GATA3 (qHsaCED0043189), ESR1 (qHsaCED0033920), TP53 (qHsaCED0045022), HK2 (qHsaCID0012715), PKM2 (qHsaCID0037934), GLS1 (qHsaCID0007574), CDH1 (qHsaCED0042076), VIM (qHsaCID0012604), and HIF1A (qHsaCED0042813). Each reaction was run in triplicate with two no-template controls per assay. Amplification data were analyzed using CFX Maestro Software (Bio-Rad). Gene expression was normalized to GAPDH and calculated using the 2⁻ΔΔCq method, with 2D untreated control as the calibrator. Three independent biological replicates were performed.

### 2.10 Statistical analysis

All quantitative data are expressed as the mean ± standard deviation. All the statistical analyses were performed in R software (R version 4.3.3). An unpaired Welch’s t-test was used for analyses and comparisons between two groups. For multiple group analyses and comparisons, one-way ANOVA and two-way ANOVA were used, with the subsequent application of Tukey’s post hoc test. IC₅₀ values were determined using four-parameter nonlinear regression. All experiments were repeated at least three times. Statistical significance was defined as*p ≤ 0.05, **p ≤ 0.01, and ***p ≤ 0.001. Principal component analysis (PCA) was performed in Julia (v1.12) on metabolic rate data after z-score normalization (autoscaling: mean-centered and scaled to unit variance per metabolite) across 13 metabolites. PCA was computed via singular value decomposition of the normalized matrix using Julia’s LinearAlgebra standard library. For each experimental group, a 95% confidence ellipse was drawn around its PC1–PC2 scores assuming bivariate normality. Pearson correlation analysis was performed separately for each treatment condition (Control and 5-FU) using Julia (v1.12). Within each condition (n = 9; three independent experiments at three time points), correlations were computed between each spheroid morphology parameter and each metabolite flux. Because these analyses were exploratory, p-values were not adjusted for multiple comparisons. Two-tailed p < 0.05 was considered statistically significant.

## 3. Results

### 3.1 5-FU suppresses spheroid growth and attenuates morphological remodeling

MCF-7 spheroids exhibited distinct morphological trajectories under control and 5-FU treatment conditions (Fig. 1A). Representative brightfield images showed that control spheroids progressively developed an irregular and elongated morphology over 72 h, whereas 5-FU-treated spheroids maintained a more spherical morphology throughout the observation period. Quantitative morphological analysis was performed using three parameters: area, aspect ratio (AR), and Feret diameter (Fig. 1B–D). Changes in spheroid shape were first assessed using AR (Fig. 1B). Control spheroids became progressively more elongated over time (p = 0.017), consistent with the microscopic observations. In contrast, AR values of 5-FU-treated spheroids remained relatively stable (1.1–1.2), indicating preservation of a near-spherical morphology. To evaluate changes in spheroid size, area and Feret diameter were normalized to baseline values at 0 h (Fig. 1C,D). 5-FU-treated spheroids exhibited a progressive reduction in size over time, with area and Feret diameter decreasing by approximately 25% (p = 0.002) and 12% (p = 0.009), respectively. In contrast, control spheroids showed an initial decrease followed by gradual increase in area, while Feret diameter exhibited similar trend as area. Taken together, untreated spheroids increased in size and progressively elongated over 72 h, whereas 5-FU treatment reduced spheroid growth and attenuated elongation.

**Fig. 1.**
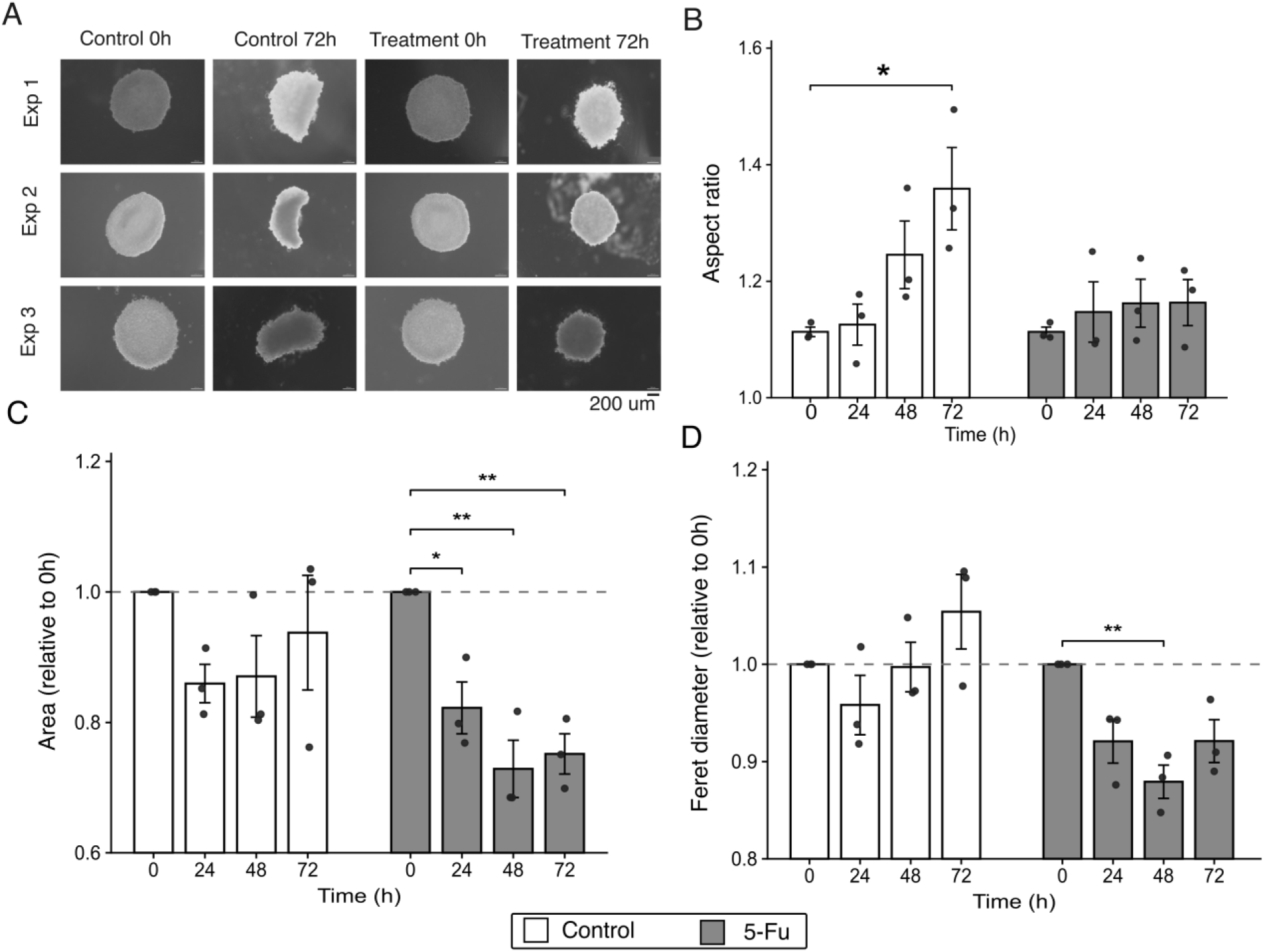
Morphological characterization of MCF-7 3D spheroids in the control and 5-FU treated sample. (A) Representative brightfield microscopy images of MCF-7 spheroids at 0 h and 72 h under control and 5-FU-treated conditions from three independent biological experiments (n = 3). Scale bar: 200 µm. (B–D) Quantification of spheroid morphology over time, including (B) aspect ratio of spheroids, (C) Normalized spheroid area and (D) normalized Feret diameter, measured at 0, 24, 48, and 72 h. Values were normalized to 0 h. Data are presented as mean ± SEM from three independent biological experiments (n = 3). Statistical significance was assessed using one-way ANOVA followed by Tukey’s HSD post hoc test. *p < 0.05, **p < 0.01.

### 3.2 Differential effects of 5-FU on viability and proliferation in 2D monolayers and 3D spheroids

To visualize and quantify the effect of 5-FU on cell viability in monolayer (2D) and spheroid (3D) cultures, we performed live/dead fluorescence staining at 0 h and 72 h. Representative fluorescence images showed that 2D control cultures maintained dense green (live) cell coverage at 72 h, while 5-FU-treated 2D cultures displayed markedly reduced live cell density compared to controls, with scattered dead (red) cells (Fig. 2A). In 3D spheroids, both control and treatment groups retained compact spheroid structures with predominantly green fluorescence at 72 h, although treated spheroids appeared slightly less bright (Fig. 2C). At 72 h, diffuse red fluorescence patches were observed in the spheroid interior. Quantification of mean green fluorescence intensity revealed a culture-dependent drug response (Fig. 2B, D). In 2D cultures, fluorescence intensity was comparable between control and 5-FU groups at 0 h. By 72 h, control cultures showed a slight increase, whereas 5-FU-treated cells decreased markedly by ∼37% (p = 0.007; Fig. 2B). In contrast, 3D spheroids showed no significant change in fluorescence intensity over time in either group, indicating a reduced response to 5-FU in the spheroid context (Fig. 2D). Overall, these results demonstrate a pronounced cytotoxic effect of 5-FU in 2D cultures, while 3D spheroids exhibit markedly attenuated drug sensitivity.

**Fig. 2.**
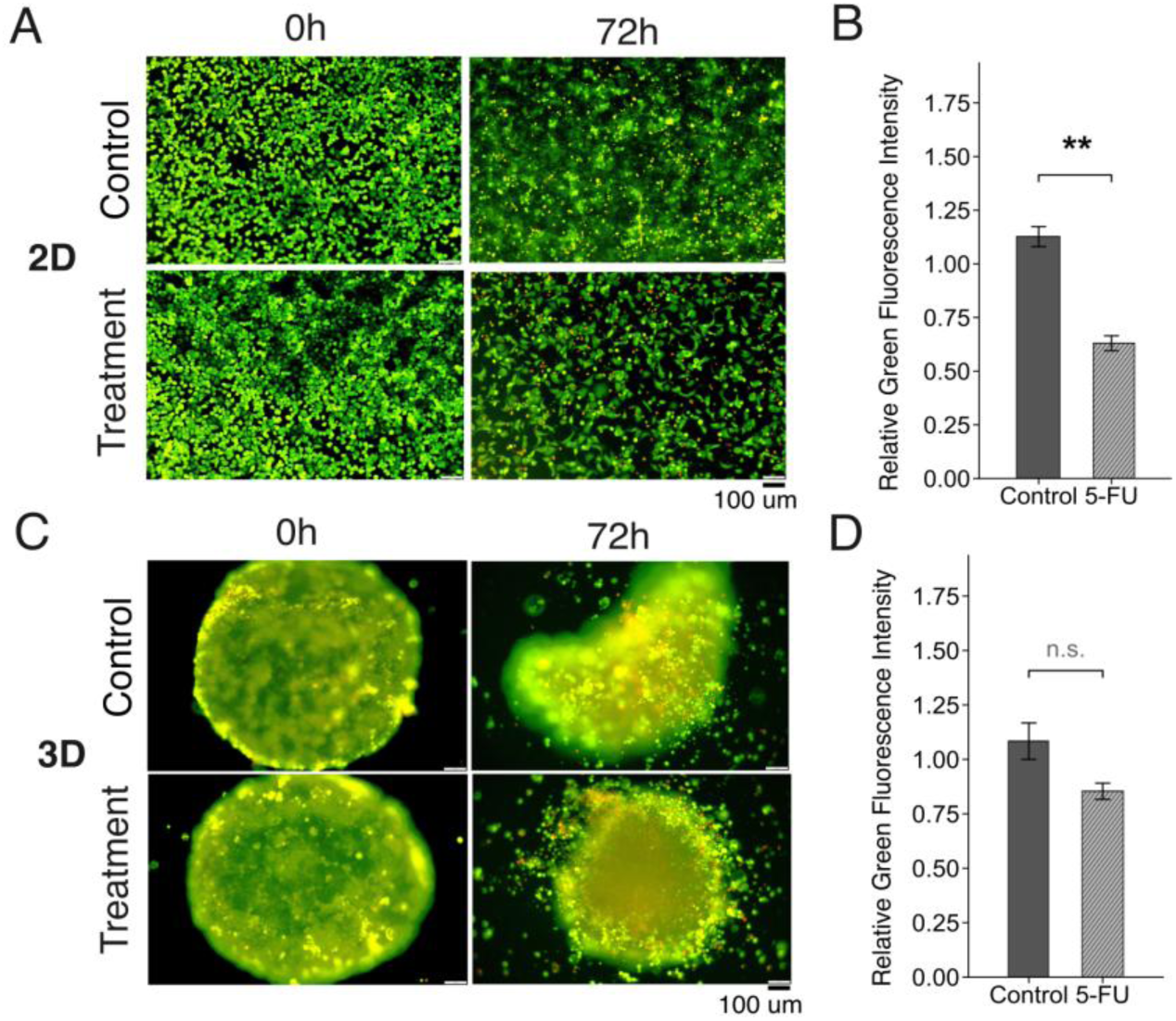
Images and quantification of LIVE/DEAD fluorescence staining of 2D monolayer and 3D spheroid cultures. Representative fluorescence microscopy images of MCF-7 cells cultured under different conditions, acquired immediately after drug addition (0 h) and after 72 h of treatment. (A) Images of 2D monolayer cultures. (B) Quantification of relative green-channel fluorescence intensity in 2D cultures, normalized to 0 h. (C) Images of 3D spheroid cultures. (D) Quantification of relative green-channel fluorescence intensity in 3D spheroids, normalized to 0 h. Scale bars = 100 µm. Data are presented as mean ± SEM from three independent biological experiments (n = 3). Statistical significance was assessed using Welch’s t-test. *p < 0.05, **p < 0.01, ns = not significant.

Viable cell numbers were calculated as the total cell count multiplied by the proportion of live cells measured by flow cytometric live/dead staining at each time point (gating strategy and live cell percentages are shown in Supplementary Fig. S2). Comparison of different cell culture models revealed distinct proliferation patterns between 2D and 3D cultures. Under control conditions, 2D monolayers increased approximately 2.7-fold over 72 h, whereas 3D spheroids maintained stable cell numbers (Fig. 3A). Two-way ANOVA revealed a significant effect of culture condition (p < 0.0005), with 2D and 3D cultures differing significantly at 72 h (Tukey HSD, p = 0.004). To further evaluate the effect of drug treatment, cell proliferation was assessed in the presence of 5-FU. In 2D cultures, viable cell numbers increased 2.67-fold over 72 h under control conditions (Fig. 3B). 5-FU treatment significantly suppressed this increase (two-way ANOVA drug effect, p = 0.003), with a 45.5% reduction in viable cell numbers compared with controls at 72 h (Tukey HSD, p = 0.02). In 3D spheroid cultures, neither the control nor the 5-FU-treated group exhibited noticeable proliferation during the 72-h observation period (Fig. 3C). Collectively, these results demonstrate that the IC₅₀ determined in 2D monolayer cultures effectively inhibited 2D cell growth, whereas the 3D spheroid structure substantially attenuated the drug response. Notably, 3D spheroids exhibited minimal proliferative activity even in the untreated control group.

**Fig. 3.**
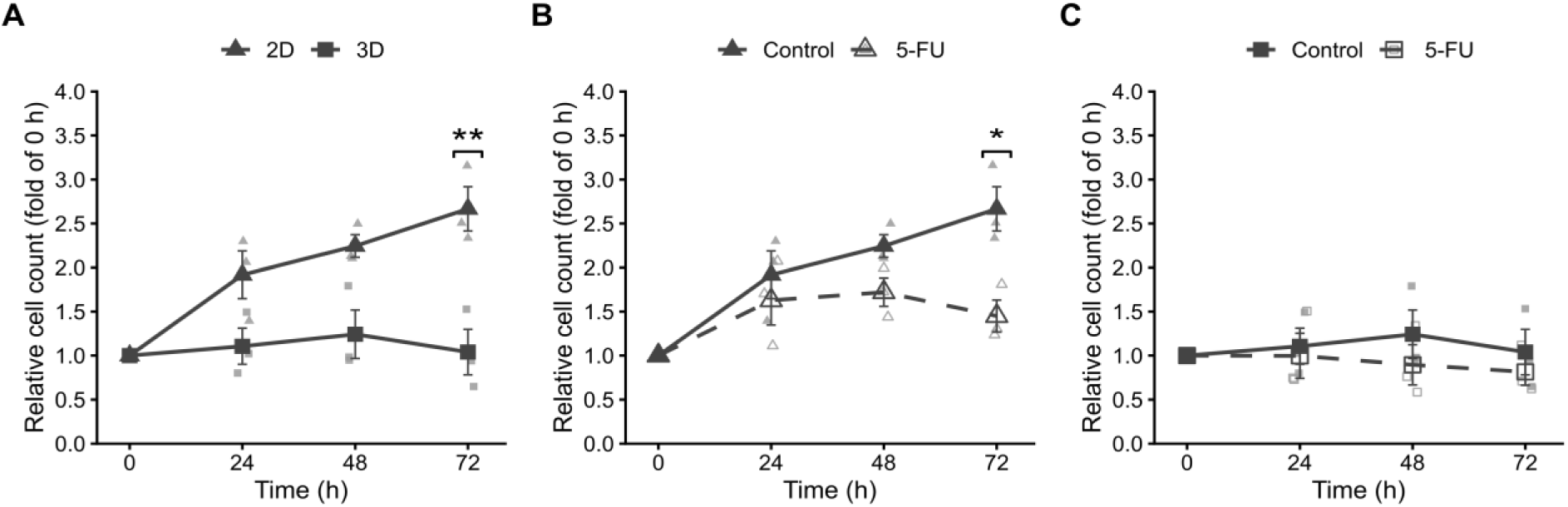
Cell proliferation in 2D monolayer and 3D spheroid cultures over time. (A) Comparison of cell proliferation between 2D and 3D culture models over 72 h. (B) Effect of 5-FU treatment on cell proliferation in 2D cultures over 72 h (2D-control vs. 2D-5FU). (C) Effect of 5-FU treatment on cell proliferation in 3D spheroid cultures over 72 h (3D-control vs. 3D-5FU). Data are presented as mean ± SEM from three independent biological experiments (n = 3). Statistical significance was assessed using two-way ANOVA followed by Tukey’s HSD post hoc test. *p < 0.05,**p < 0.01.

### 3.3 3D spheroids exhibit increased GLUT1 positive population compared with 2D monolayers

Given the role of GLUT1 in regulating glucose uptake and glycolytic activity in cancer cells (Szablewski, 2013), we assessed whether culture architecture affected GLUT1-positive cell populations. Flow cytometric analysis revealed distinct GLUT1-positive populations between 2D monolayers and 3D spheroids (Fig. 4A–D). Representative histograms at 72 h showed a higher GLUT1-positive population in 3D spheroids compared with 2D monolayers (Fig. 4A). Quantification across the 72 h observation period confirmed a consistently greater fraction of GLUT1-positive cells in 3D spheroids (Fig. 4B). At 24 h, the proportion of GLUT1-positive cells in 3D control spheroids was approximately 5.8-fold higher than that in 2D controls, and this increase was maintained at 48 h and 72 h. Two-way ANOVA revealed a significant effect of culture condition on the GLUT1-positive cell population (p < 0.001), with Tukey post hoc analysis confirming significant differences between 2D and 3D cultures at 24 h (p = 0.003) and 72 h (p = 0.038). Furthermore, 5-FU treatment did not significantly affect the GLUT1-positive cell population in either culture model. No significant drug effect was observed in either 2D monolayers or 3D spheroids (Fig. 4C,D). Overall, the proportion of GLUT1-positive cells was consistently higher in 3D spheroids than in 2D monolayers throughout the observation period. Although 5-FU treatment caused minor variations in the GLUT1-positive population, culture architecture was the primary factor associated with GLUT1 differences, rather than drug exposure.

**Fig. 4.**
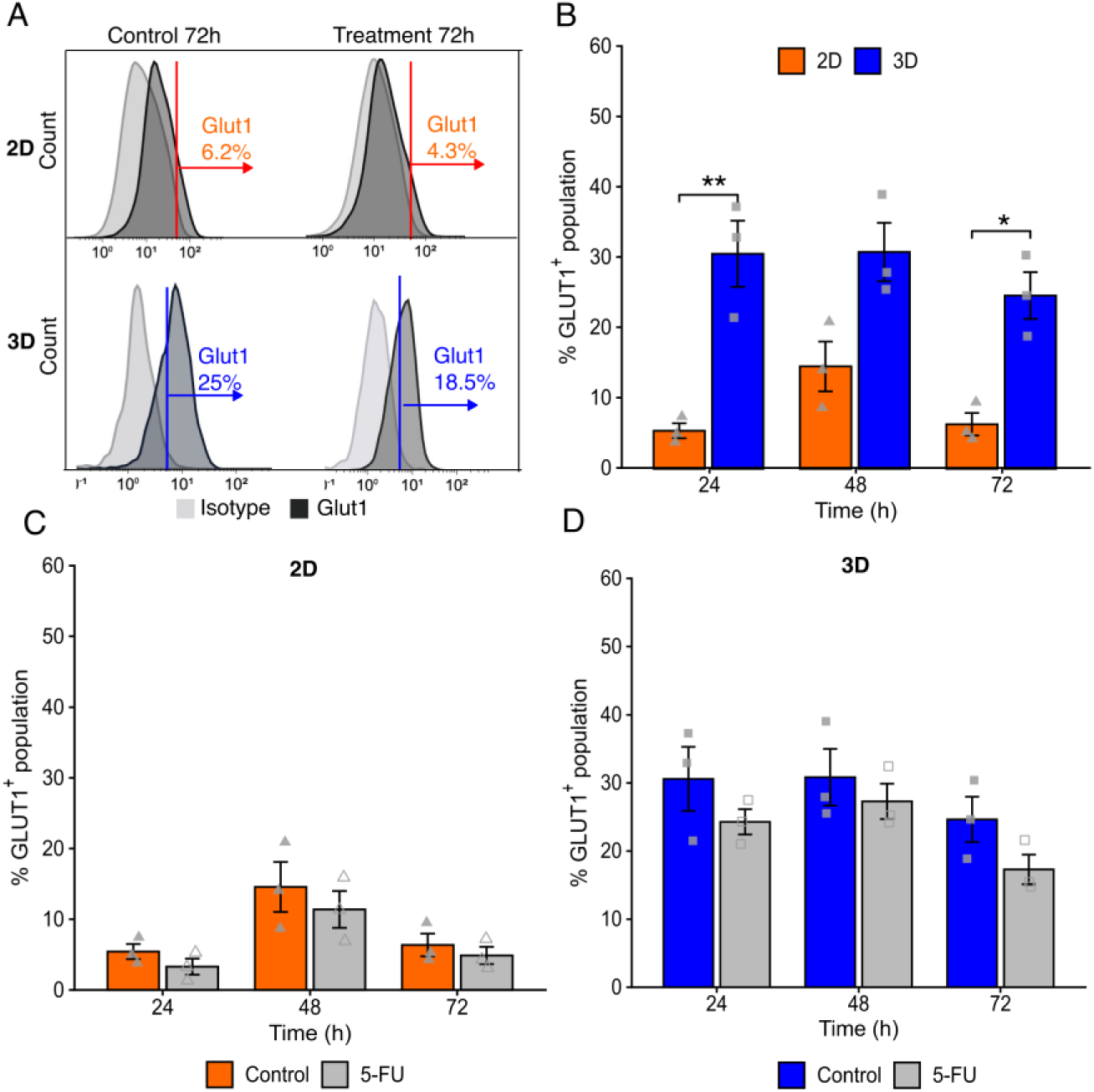
GLUT1^+^ population in 2D monolayer and 3D spheroid cultures assessed by flow cytometry. (A) Representative flow cytometry histograms of GLUT1^+^ population versus isotype control (blue) at 72 h in 2D (top) and 3D (bottom) cultures in control (left) and 5-FU-treated (right) conditions. Percentages indicate GLUT1^+^ cell populations. (B) Quantification of GLUT1^+^ cell percentage in 2D and 3D control group at 24, 48, and 72 h. (C) GLUT1^+^ cell percentage in 2D monolayer over time in the control and 5-FU-treated conditions. (D) GLUT1^+^ cell percentage in 3D spheroids over time in the control and 5-FU-treated conditions. Data are presented as mean ± SEM from three independent biological experiments (n = 3). Statistical significance was assessed using two-way ANOVA followed by Tukey’s HSD post hoc test. *p < 0.05,**p < 0.01.

### 3.4 Distinct metabolic flux dynamics between 2D and 3D cultures under 5-FU treatment

To determine whether culture-dependent differences in glucose uptake capacity were reflected in metabolic activity, we next quantified extracellular metabolic fluxes across 2D monolayers and 3D spheroids under control and 5-FU-treated conditions. Fluxes were normalized to viable cell number(fmol/cell/h; see Methods), allowing comparison of metabolic activity on a per-viable-cell basis across culture conditions. Additional metabolite flux profiles are shown in Supplementary Fig. S3.

Glucose consumption and lactate production displayed distinct temporal patterns and drug responses between 2D and 3D cultures (Fig. 5A,B). In 2D monolayers, per-viable-cell glucose consumption and lactate production gradually declined over the 72 h observation period under control conditions, with both rates decreasing by approximately 29% from 24 to 72 h. 5-FU treatment produced a modest reduction in glucose consumption and lactate production at 48–72 h (∼15% and ∼18%, respectively), with minimal effects at 24 h. In contrast, 3D spheroids displayed a distinct temporal pattern. Although initially lower than that in 2D monolayers at 24 h, glucose consumption in 3D spheroids increased over time, rising by approximately 23% from 24 to 72 h.

**Fig. 5.**
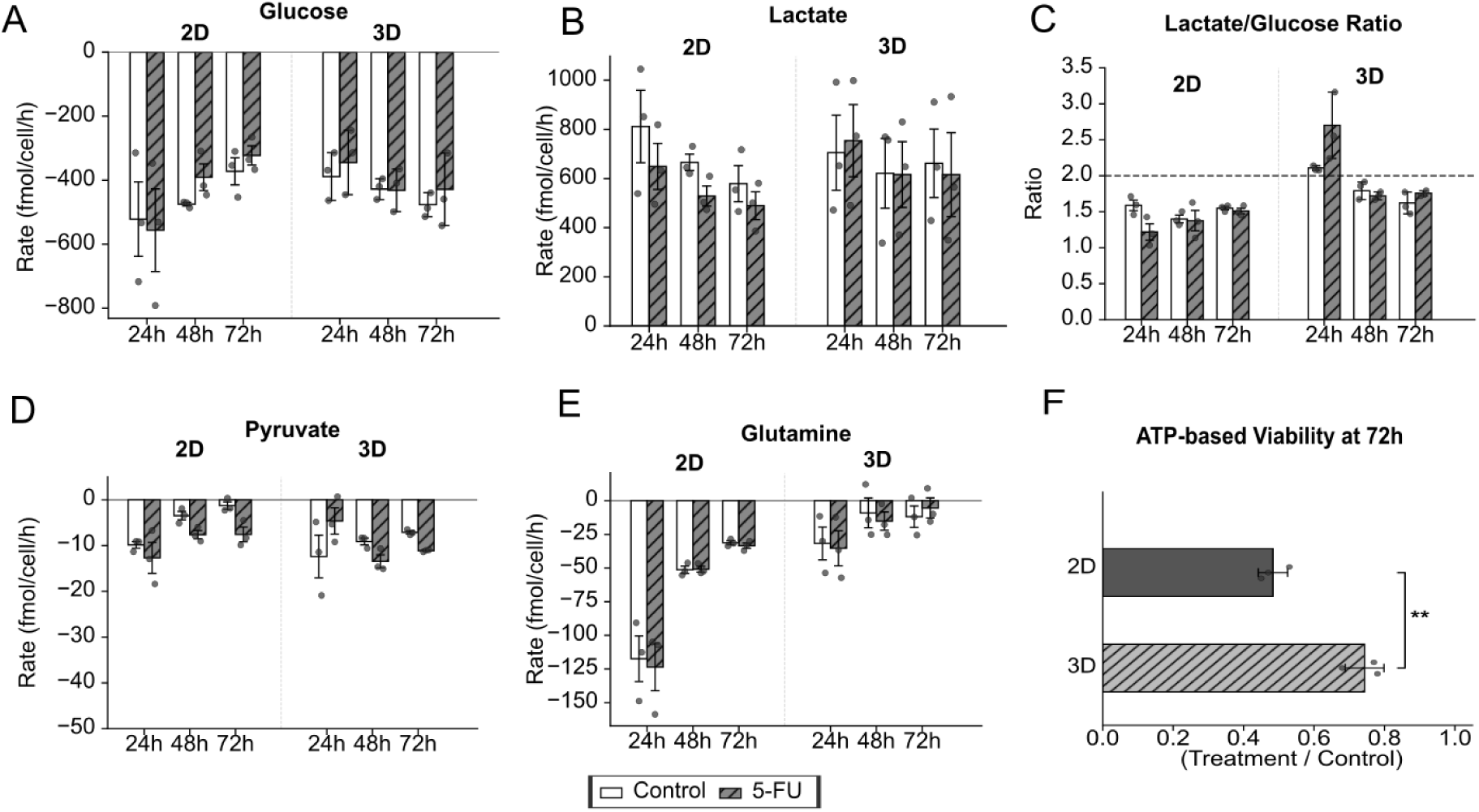
Metabolic rates and energy metabolism in 2D monolayer and 3D spheroid cultures. Metabolic consumption and production rates were quantified by ¹H NMR exometabolomics at 24, 48, and 72 h and normalized to viable cell number. (A–E) Metabolic fluxes in 2D (left) and 3D (right) cultures. Open bars, control; hatched bars, 5-FU-treated. (A) Glucose consumption rate. (B) Lactate production rate. (C) Lactate-to-glucose ratio (glycolytic index); the dashed line indicates the theoretical maximum value (2.0) for complete glycolytic conversion of glucose to lactate. (D) Pyruvate consumption rate. (E) Glutamine consumption rate. (F) Relative ATP-based viability at 72 h, expressed as ATP signal in 5-FU-treated cultures relative to untreated controls. Data are presented as mean ± SEM from three independent biological experiments (n = 3) for each culture model. Statistical significance was assessed using two-way ANOVA followed by Tukey’s HSD post hoc test for panels A–E, and Welch’s *t*-test for panel F. \**p* < 0.05, \*\**p* < 0.01.

Lactate production in control 3D spheroids remained relatively stable throughout the observation period, and 5-FU treatment caused only minor changes in glucose consumption and lactate production fluxes. Consistent with these differences, the lactate-to-glucose ratio (L/G ratio; Fig. 5C) was consistently higher in 3D spheroids than in 2D monolayers across the observation period.

Pyruvate uptake showed distinct temporal dynamics between 2D and 3D cultures (Fig. 5D). Under control conditions, pyruvate uptake declined markedly in 2D monolayers over time, decreasing by approximately 87% from 24 to 72 h. In contrast, 3D spheroids showed a much less pronounced decrease, with uptake remaining relatively stable throughout the observation period. 5-FU treatment increased pyruvate uptake in both culture models, particularly at later time points, although the absolute flux values remained low and should therefore be interpreted cautiously.

Glutamine consumption showed distinct culture-dependent patterns (Fig. 5E). In 2D monolayers, glutamine consumption declined progressively over time, decreasing by approximately 73% from 24 to 72 h. In contrast, 3D spheroids exhibited consistently lower glutamine consumption than 2D cultures (p < 0.001) and maintained relatively stable rates over time. 5-FU treatment did not significantly alter glutamine consumption in either culture model. Glutamate release showed a similar culture-dependent pattern (Supplementary Fig. S3), with higher levels in 2D cultures than in 3D spheroids. While glutamate release declined over time in 2D cultures, 3D spheroids maintained relatively stable levels. In both culture models, glutamate release remained lower than glutamine consumption, although both fluxes were within the same order of magnitude.

For the remaining metabolites (Supplementary Fig. S3), flux profiles were largely comparable between 2D and 3D cultures, with only minor temporal differences observed. 5-FU treatment had minimal effects, and none of the differences reached statistical significance.

Relative ATP activity was assessed at 72 h to evaluate the impact of 5-FU on cellular energy status (Fig. 5F). In 2D monolayers, 5-FU treatment reduced ATP activity by 52% compared with untreated controls. In contrast, 3D spheroids maintained significantly higher ATP activity following 5-FU exposure (p = 0.004), with only a 26% reduction relative to controls.

Overall, metabolic flux profiling revealed distinct temporal trajectories between 2D and 3D cultures. 3D spheroids showed increased metabolic flux over time compared with 2D monolayers, while 5-FU treatment caused limited changes in metabolite fluxes and ATP activity.

### 3.5 Culture architecture shapes metabolic variation and morphology-associated single-cell metabolic flux in 3D spheroids

Given the distinct metabolic profiles observed between 2D and 3D cultures, we next performed principal component analysis (PCA) to evaluate the major sources of metabolic variation across experimental conditions. PCA was performed using the 13 NMR-quantified metabolite exchange rates from all experimental groups (Fig. 6A). The first two principal components accounted for 66.0% of the total variance (PC1: 38.9%; PC2: 27.1%). Samples were primarily separated along PC1 according to culture model, with 2D and 3D cultures occupying distinct score regions, whereas separation between control and 5-FU-treated groups was less pronounced. The loading plot (Fig. 6B and S4) indicated that PC1 was driven by multiple metabolite exchange rates, including glycolytic metabolites (glucose, pyruvate, and lactate) as well as amino acid-related metabolites (glutamine, glutamate, BCAAs, and aromatic amino acids). PC2 was associated with temporal variation among samples. Together, these results identified culture model as the dominant source of metabolic variation.

**Fig. 6.**
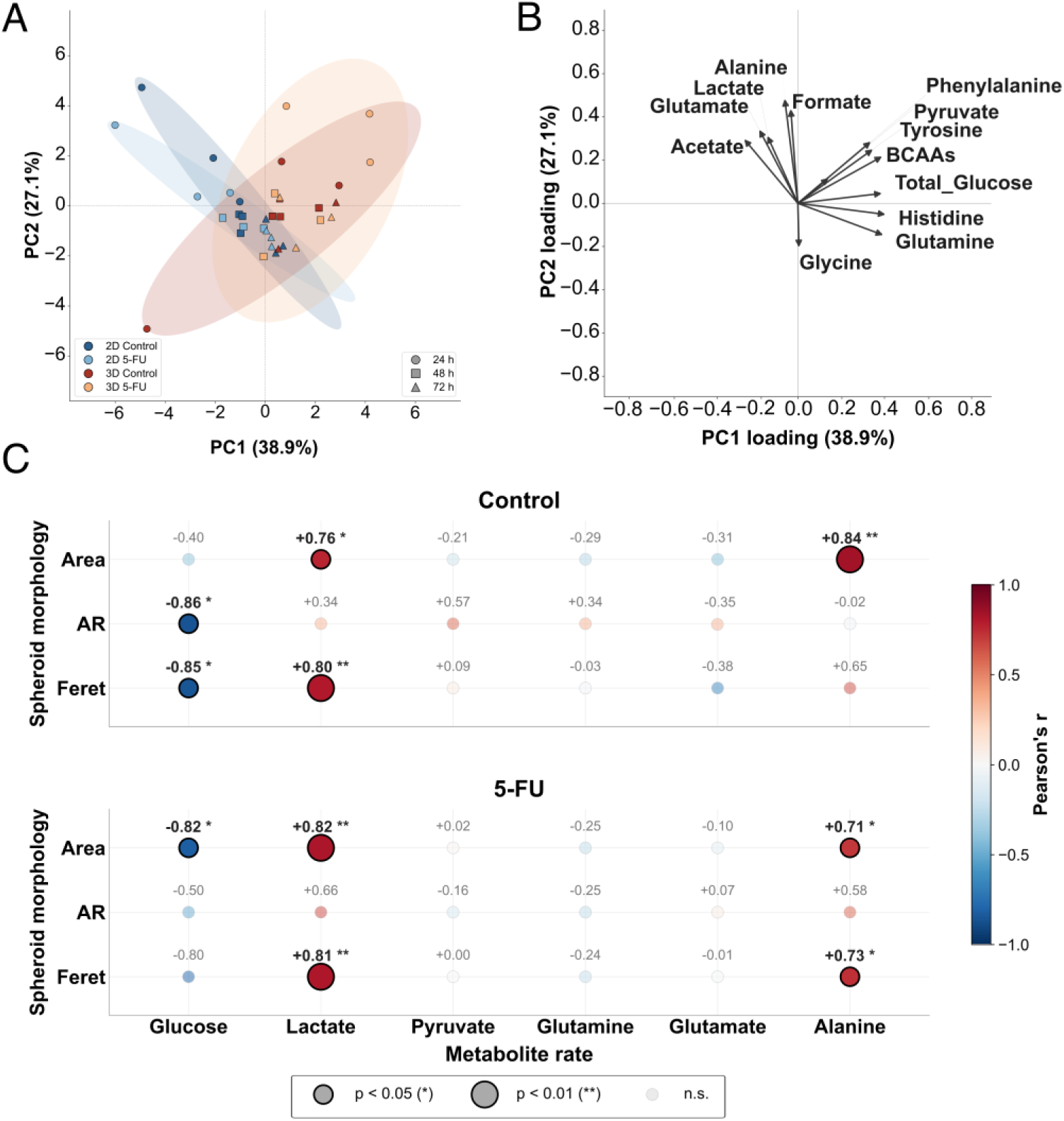
Integrated multivariate analysis of metabolic rates and morphology–metabolism correlations. Principal component analysis (PCA) of metabolic rates (A), and Pearson correlation waterfall plots of morphology–metabolite associations in control (B) and 5-FU-treated (C) spheroids. Shaded ellipses indicate 95% confidence intervals. Data were z-score normalized prior to PCA; loadings are shown in Supplementary Fig. S4. Each point represents a morphology– metabolite pair (n = 9 per condition). Point size reflects statistical significance, and colour indicates the direction of correlation. Only correlations with *p* < 0.05 are labelled.

We next investigated whether spheroid morphology contributed to the metabolic differences observed within 3D cultures. Correlation analysis between spheroid morphological parameters and metabolic fluxes was performed across 3D culture conditions (Fig. 6C). Because each flux was normalized to viable cell number, these correlations reflect morphology-associated differences in metabolic activity at the level of individual viable cells. Under both control and 5-FU-treated conditions, spheroid size (area and Feret diameter) was positively correlated with lactate and alanine release, whereas glucose flux was negatively correlated with spheroid size, indicating that larger spheroids were associated with increased glucose consumption and lactate/alanine production on a per-cell basis. Moreover, AR was negatively correlated with glucose flux, suggesting that more elongated spheroids were associated with increased per-cell glucose consumption. Overall, spheroid morphology was associated with per-viable-cell metabolic flux, suggesting that metabolic heterogeneity within 3D cultures is linked to differences in spheroid architecture.

### 3.6 Gene expression profiling reveals culture-dependent glycolytic reprogramming and epithelial phenotype reinforcement

To determine whether the culture-dependent differences in GLUT1 expression and metabolic flux were accompanied by transcriptional changes, quantitative RT–PCR analysis of genes associated with cellular phenotype and metabolism was performed at the 72 h endpoint (Fig. 7).

**Fig. 7.**
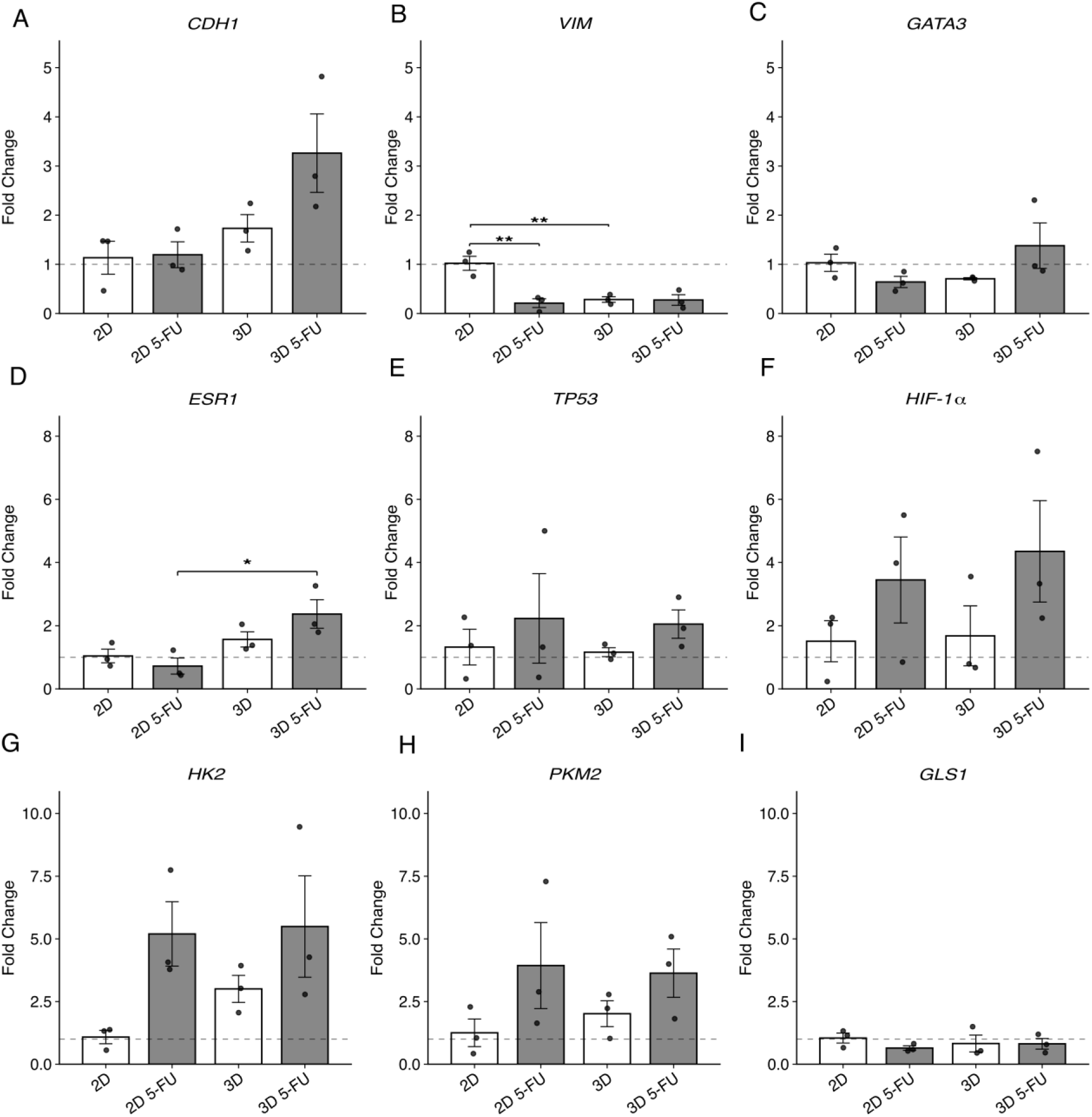
Gene expression profiling by quantitative RT-PCR at 72. **h.** Relative mRNA expression of eight target genes. (A) CDH1, (B) VIM, (C) GATA3, (D) ESR1, (E) TP53, (F) HIF-1α, (G) HK2, (H) PKM2, (I) GLS1. Bars represent mean fold change ± SEM from three biological replicates (n = 3); individual data points are shown as black dots. The dashed horizontal line at fold change = 1 indicates the 2D-Control reference level. Statistical significance was assessed using one-way ANOVA followed by Tukey’s HSD post hoc test across the four groups. Only statistically significant pairwise comparisons (p < 0.05) are shown. *p < 0.05, **p < 0.01.

Among epithelial–mesenchymal transition (EMT)-related markers, the 3D culture model displayed a more epithelial-like transcriptional profile than 2D monolayers. Under untreated conditions, VIM expression was significantly lower in 3D spheroids than in 2D controls (p = 0.0045), whereas CDH1 expression was moderately increased. Upon 5-FU treatment, VIM expression decreased further in 2D cultures but remained stable in 3D spheroids; conversely, CDH1 expression was further elevated after drug exposure in 3D spheroids. These findings demonstrate a reciprocal regulation of CDH1 and VIM across culture systems and treatments. Among lineage-associated genes, ESR1 showed the most pronounced response, with transcript levels significantly higher in 5-FU treated 3D spheroids than in the corresponding 2D cultures (p = 0.021). GATA3 and TP53 showed modest increases after 5-FU treatment in both culture systems, but differences between groups were not significant. Within the metabolic gene panel, HK2 and PKM2 were elevated in untreated 3D cultures relative to 2D cultures and increased further after 5-FU exposure; HIF1A showed a similar upward trend in both treatment groups, while GLS1 remained largely unchanged. None of these metabolic gene changes reached statistical significance

In short, qPCR profiling confirmed that 3D spheroids maintained a transcriptional profile characterized by elevated CDH1, ESR1, and glycolytic gene expression relative to 2D monolayers, a trend that was further pronounced upon 5-FU treatment.

## 4. Discussion

Pharmacological responses characterised in 2D monolayers frequently fail to translate to human tumours (Hammond *et al*, 2024), motivating the migration to 3D models (Jensen & Teng, 2020); yet how culture architecture jointly reshapes metabolism and cellular behavior under therapeutic stress remains poorly defined. Here, we demonstrate that culture dimensionality, rather than drug exposure, is the dominant determinant of the metabolic phenotype in MCF-7 cells. By combining time-resolved ¹H NMR cellular metabolomics with morphological, transcriptional, and viability readouts in MCF-7 monolayers and spheroids exposed to 5-FU, we show that 3D architecture coordinates proliferation, epithelial organization, and glycolytic activity into a structure-defined metabolic state. Within this framework, 5-FU functions as a secondary perturbation. This perspective reorients the interpretation of in vitro drug responses, establishing that apparent pharmacological effects must be evaluated against the baseline metabolic state imposed by the culture model itself.

Morphological and transcriptional phenotypes also diverged from 2D in ways congruent with architecture-defined regulation. 3D culture provides more extensive cell–cell interactions (Mangani *et al*, 2025). We found the untreated spheroids progressively elongated over 72 h, whereas 5-FU-treated spheroids remained compact and near-spherical. One possible explanation is that 5-FU preferentially targets proliferating cells (Hirschhaeuser *et al*, 2010). Thus, the inhibition of proliferative activity may reduce the dynamic cellular rearrangements required for spheroid elongation—a process also indicative of drug-induced changes in cell-cell contacts within the spheroid. E-cadherin-mediated adherens junctions maintain epithelial tissue integrity (Coopman & Djiane, 2016), and previous studies have linked 3D culture conditions to increased CDH1 expression and enhanced spheroid compactness (Smyrek *et al*, 2019). At the transcriptional level, we found 3D culture reinforced an epithelial signature, with elevated CDH1 and reduced VIM expression, and 5-FU further amplified this pattern, which suggests strengthened intercellular adhesion, which may explain why spheroids in the drug-treated group gradually shrank while consistently maintaining a spherical shape. Together, structural and transcriptional readouts suggest that 3D architecture both restrains growth and consolidates a stress-resistant epithelial state.

Within this structure-defined state, the most pronounced divergence between 2D and 3D cultures emerged in glycolytic regulation. In breast cancer, elevated GLUT1 and HK2 expression is commonly associated with increased glycolytic activity (Garg *et al*, 2025; Wu *et al*, 2025; Liu *et al*, 2024), and 3D culture systems have previously been reported to exhibit enhanced glycolytic activity compared with 2D monolayers (Takakusagi *et al*, 2025; Al-Masri *et al*, 2021). Our study further focused on per-cell metabolic activity under different culture conditions. Over the 72 h treatment period, per-cell glucose uptake decreased in 2D cultures but increased in 3D spheroids, suggesting that cells in 3D developed a more glycolytic metabolic state. This was further supported by the higher L/G ratio observed in 3D cultures throughout the experiment. This trajectory indicates that 3D architecture progressively reinforces glycolytic capacity, a temporal signature endpoint sampling alone would have missed. Meanwhile, the GLUT1-positive population expanded markedly, and HK2 transcript was elevated relative to monolayers, in line with prior reports of 3D-induced glycolytic shifts in other tumour models (Wang *et al*, 2021; Tidwell *et al*, 2022). Notably, in a microfluidic NMR chamber, MCF-7 spheroids consumed glucose and produced lactate lower rates than monolayer cells under closed conditions (Patra *et al*, 2021). One explanation is that maybe the metabolic behavior of spheroids is influenced by their microenvironment. In microfluidic chambers, limited oxygen and nutrient diffusion for cells at the spheroid (Grimes *et al*, 2014; Mueller-Klieser *et al*, 1986), thereby reducing the overall metabolic flux. However, 5-FU produced only lightly changes in glycolytic flux. Interestingly, this glycolytic configuration may was associated with markedly higher ATP preservation under 5-FU in 3D than in 2D, despite similar viable cell number. This indicates that the drug’s perturbation of cellular metabolic behaviour was less significant than the effects of establishing the cellular model and microenvironment itself. Considering that the oxygen- and nutrient-restricted regions inherent to multicellular spheroids constrain oxidative phosphorylation and thereby favour glycolytic ATP production (Riffle & Hegde, 2017), improved energy homeostasis under stress would be an expected consequence. This metabolic adaptation may explain why 3D spheroids tolerate 5-FU treatment more effectively than monolayer cultures.

Amino acid metabolism diverged along a separate axis. Glutamine consumption and glutamate release were both substantially lower in 3D spheroids than in 2D monolayers, with 5-FU producing no consistent effect in either format. Reduced glutamine utilisation accords with the broader observation that luminal A MCF-7 cells display weaker glutamine dependence than the canonical “glutamine-addicted” tumour phenotype (Li *et al*, 2023; Damiani *et al*, 2017), and is further amplified in the present case by the diffusion- and accessibility-limited interior of spheroids (DeBerardinis & Cheng, 2010). This discordance has practical implications: glutaminase- and amino acid–targeted therapies developed against glutamine-addicted models in 2D may be poorly aligned with the actual flux landscape of luminal breast spheroids. It therefore underscores a recurrent theme of this work—metabolic vulnerabilities inferred from monolayer culture are not necessarily portable across architectures, and culture-specific flux profiling is required to identify therapeutic entry points relevant to the in vivo geometry (Rogers *et al*, 2023).

Building on the observation that cells cultured under different culture conditions exhibit distinct metabolic behaviors, we found that both spheroid size and shape were correlated with the metabolic rates of individual cells. In both the treatment and control groups, cells within larger spheroids exhibited higher glucose consumption rates as well as higher lactate and alanine production rates on per viable cell, indicate that the higher glycolytic activity observed in larger spheroids cannot be explained by their larger size and greater cell number, but is associated with increased metabolic activity on a per viable cell. The increase in single-cell glycolytic flux may represent a metabolic consequence of microenvironmental changes that develop during spheroid growth within larger three-dimensional structures. Larger spheroids are expected to generate steeper oxygen and nutrient gradients (Hirschhaeuser *et al*, 2010). Together with the enhanced cell–cell interactions observed in 3D culture, these architectural features may contribute to metabolic adaptation at the level of individual cells. We further found that spheroid elongation was correlated with cellular metabolism, providing additional evidence that the three-dimensional architecture itself influences cellular metabolism. The observed association may reflect differences in the internal microenvironment accompanying spheroid elongation, such as altered mechanical stress (Helmlinger *et al*, 1997), nutrient availability (Park *et al*, 2016), thereby influencing the metabolic behavior of the constituent cells. Although these mechanistic explanations remain preliminary, our findings indicate that spheroid morphology and structure further influence metabolic adaptation at the level of individual viable cells. This offers valuable insights for the future development and selection of drug-response models.

As the analysis was carried out in a single luminal A (ER⁺) model, the conclusions drawn here pertain primarily to this molecular subtype and are not directly extrapolated to other breast cancer phenotypes; this is particularly pertinent for more metabolically distinct subtypes such as triple-negative breast cancer cells, which exhibit substantially different metabolic programmes (Sun *et al*, 2020). Within the present model, metabolic rates also varied dynamically across the culture period, and discrete time-point sampling resolves only the principal trajectory of this adaptation. Real-time monitoring of metabolic fluxes—for instance through integration with microfluidic NMR platforms developed by Utz and colleagues (Patra *et al*, 2021; Barker *et al*, 2026)—would provide the higher temporal and spatial resolution required to capture dynamic metabolic reprogramming in 3D more fully. Within these scope considerations, the integrated multi-level dataset presented here quantifies how 3D architecture reshapes both the metabolic phenotype of MCF-7 cells and their response to chemotherapeutic challenge, underscoring the value of 3D models in the interpretation of preclinical drug responses and providing a quantitative reference for studies that aim to relate in vitro metabolic measurements to the metabolic environment of solid tumors.

## 5. Conclusions

This study investigated how 3D culture architecture alters the cellular metabolic state and 5-FU response of MCF-7 luminal breast cancer cells relative to conventional 2D monolayer culture. Compared with 2D cultures, 3D spheroids progressively developed a more glycolytic metabolic state at the per-cell level, characterized by a higher GLUT1-positive cell population, increased expression of metabolism-related genes, and better preservation of ATP levels. Furthermore, changes in spheroid morphology were associated with differences in per-cell glycolytic activity.

Overall, culture dimensionality emerged as a stronger determinant of cellular behavior and metabolic state than drug treatment. These results highlight that model selection substantially influences the apparent efficacy of therapeutic agents, underscoring the importance of developing physiologically relevant in vitro models that better recapitulate the in vivo microenvironment.

## Acknowledgements

This work was supported by the Helmholtz Association, the Otto-Lehmann-Professorship of M.U. at the Karlsruhe Institute of Technology (KIT), and a doctoral fellowship to Y.L. from the China Scholarship Council (CSC).

## Author Contributions

**Y.L.**: Methodology, Investigation, Formal analysis, Data curation, Visualization, Writing – original draft. **N.S.**: Methodology. **S.B.**: Writing – review & editing. **M.U.**: Conceptualization, Methodology, Software, Resources, Funding acquisition, Project administration, Writing – review & editing. **A.R.**: Conceptualization, Supervision, Project administration, Writing – review & editing.

## Data Availability

The ¹H NMR raw spectra and processed metabolite concentration tables generated and analysed in this study are available from the corresponding author upon reasonable request. Source data for all main figures are provided as supplementary files.

## Supplementary Figure Legends

**Supplementary Fig. S1.**
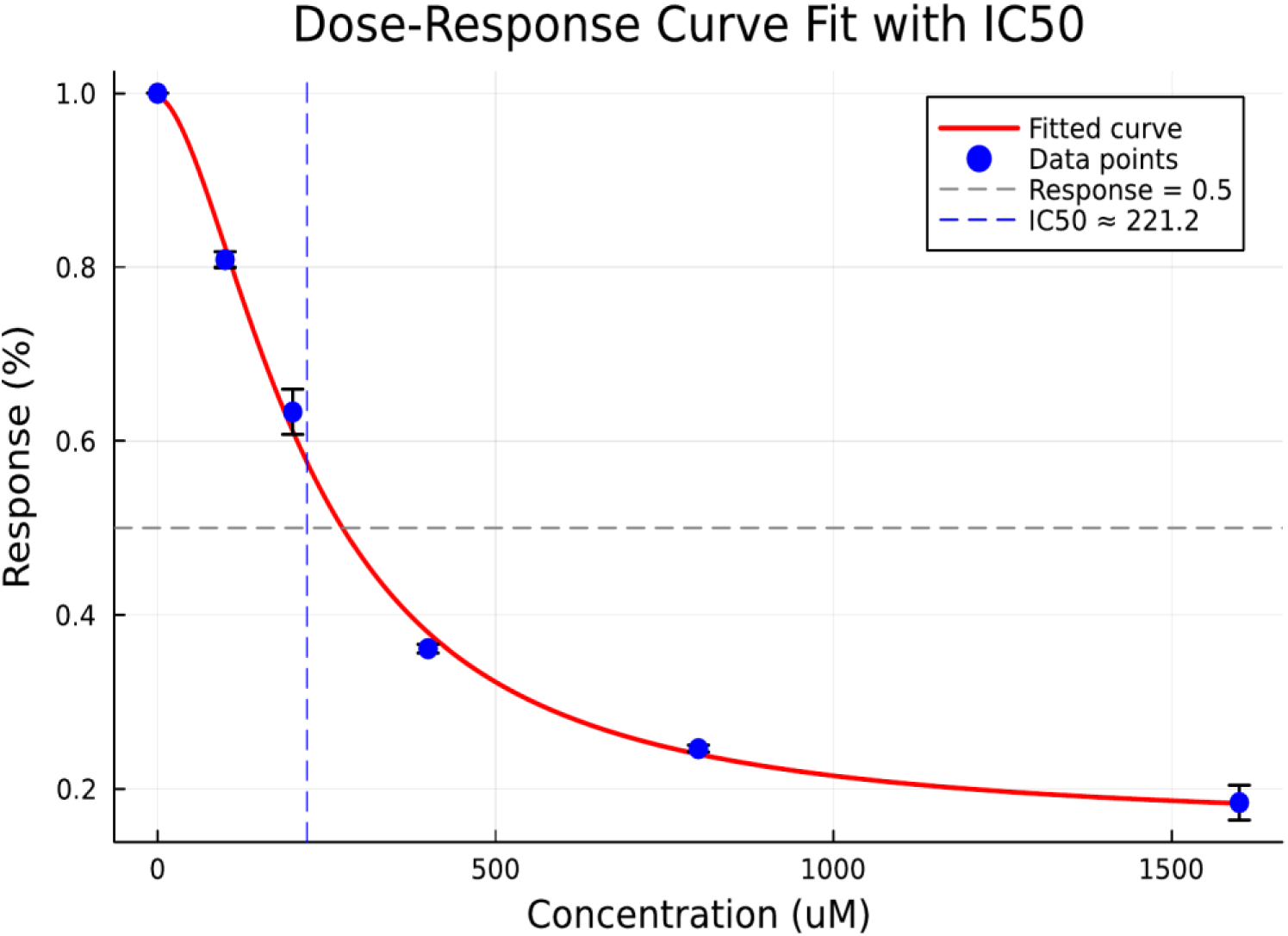
IC₅₀ determination of 5-FU in 2D MCF-7 monolayer cultures. Dose–response curve showing the effect of 5-FU (0–1600 µM) on MCF-7 cell viability after 72 h treatment, measured by CellTiter-Glo luminescence assay. Viability is expressed as a percentage relative to untreated controls. The IC₅₀ was determined by nonlinear regression (four-parameter logistic fit) to be approximately 220 µM. Data are presented as mean ± SEM (n = 3 independent experiments).

**Supplementary Fig. S2.**
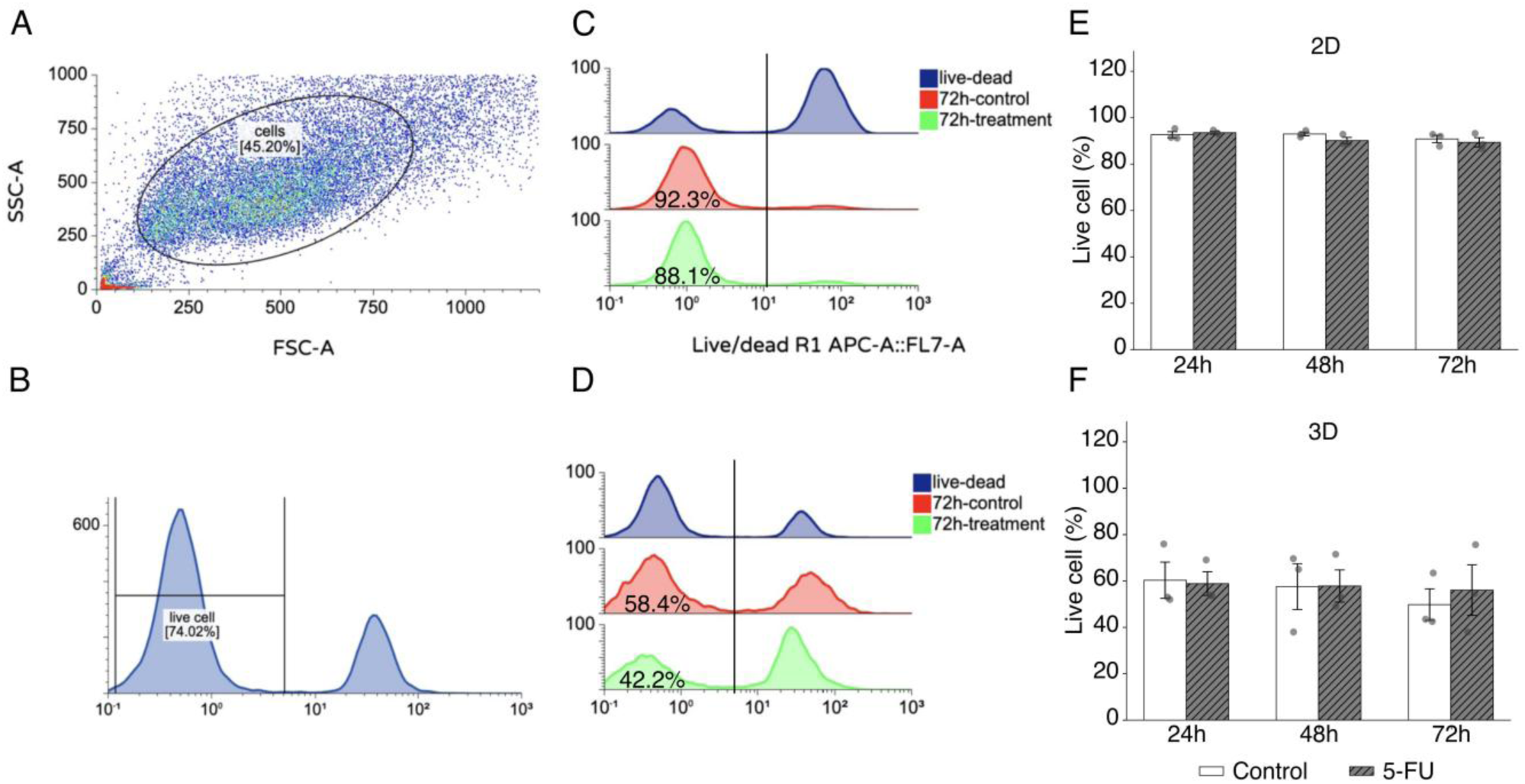
Flow cytometry gating strategy and live cell percentages. (A) Representative FSC-A versus SSC-A scatter plot showing the cell population gate used for downstream analysis. (B) Representative live/dead fluorescence histogram of dissociated 3D spheroid cells stained with LIVE/DEAD Fixable Far-Red Dead Cell Stain, illustrating the gating threshold for live cell discrimination. (C, D) Representative overlay histograms of live/dead staining for 2D cultures (C) and 3D spheroids (D) at 72 h. (E, F) Quantification of live cell percentages in 2D cultures (E) and 3D spheroids (F) at 24, 48, and 72 h. Open bars represent control and hatched bars represent 5-FU-treated samples. Data are presented as mean ± SEM (n = 3); individual data points are shown.

**Supplementary Fig. S3.**
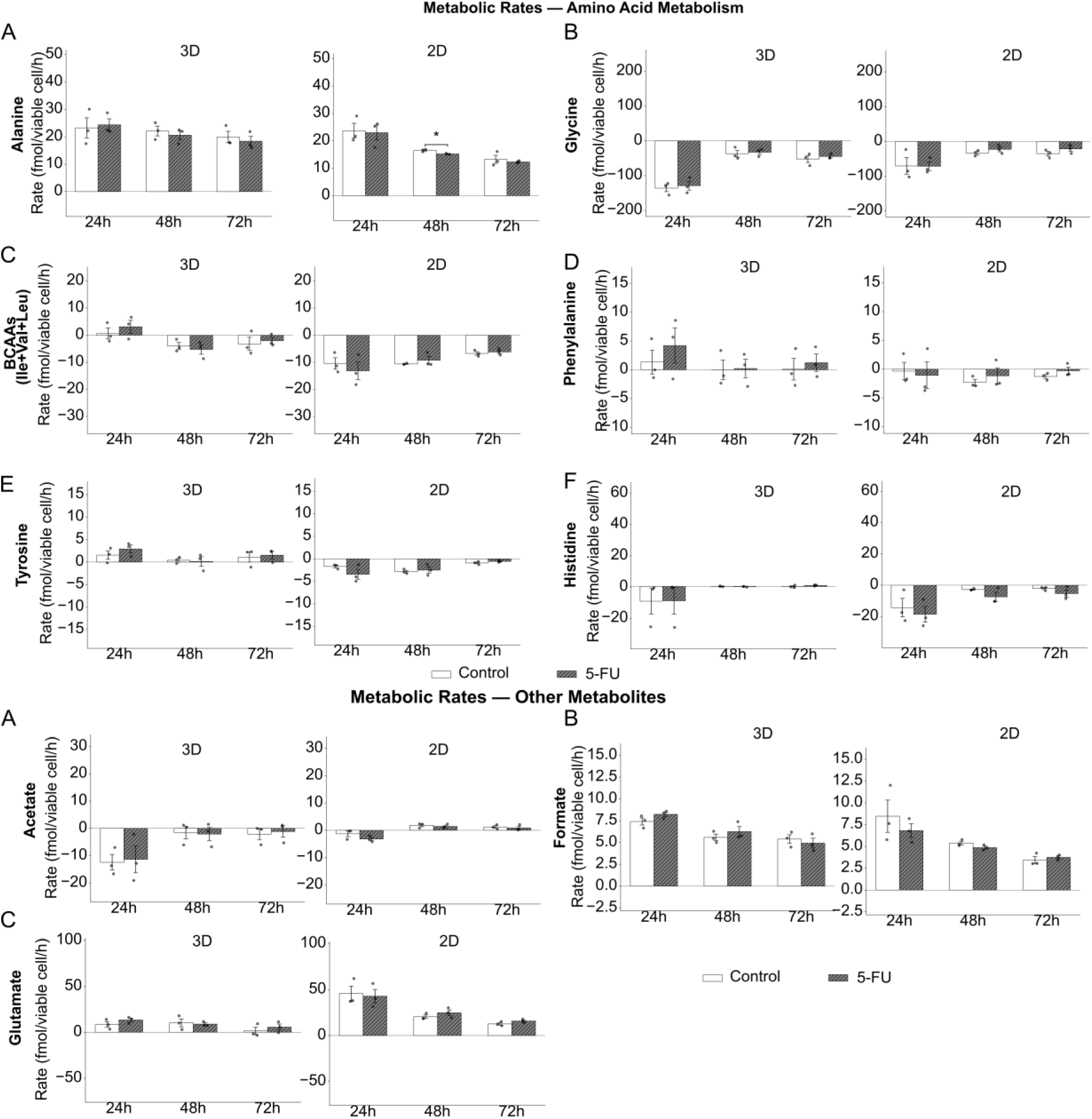
Time-resolved metabolic rates of the remaining 9 metabolites. Time-resolved metabolic rates (fmol/cell/h) for glutamate, alanine, glycine, branched-chain amino acids (BCAAs: isoleucine + valine + leucine), phenylalanine, tyrosine, histidine, acetate, and formate in 2D (left) and 3D (right) cultures at 24, 48, and 72 h. Open bars: control; hatched bars: 5-FU. Data are presented as mean ± SEM (n = 3); individual data points are shown. *p < 0.05 (Welch’s t-test, control vs. 5-FU at individual timepoints).

**Supplementary Fig. S4.**
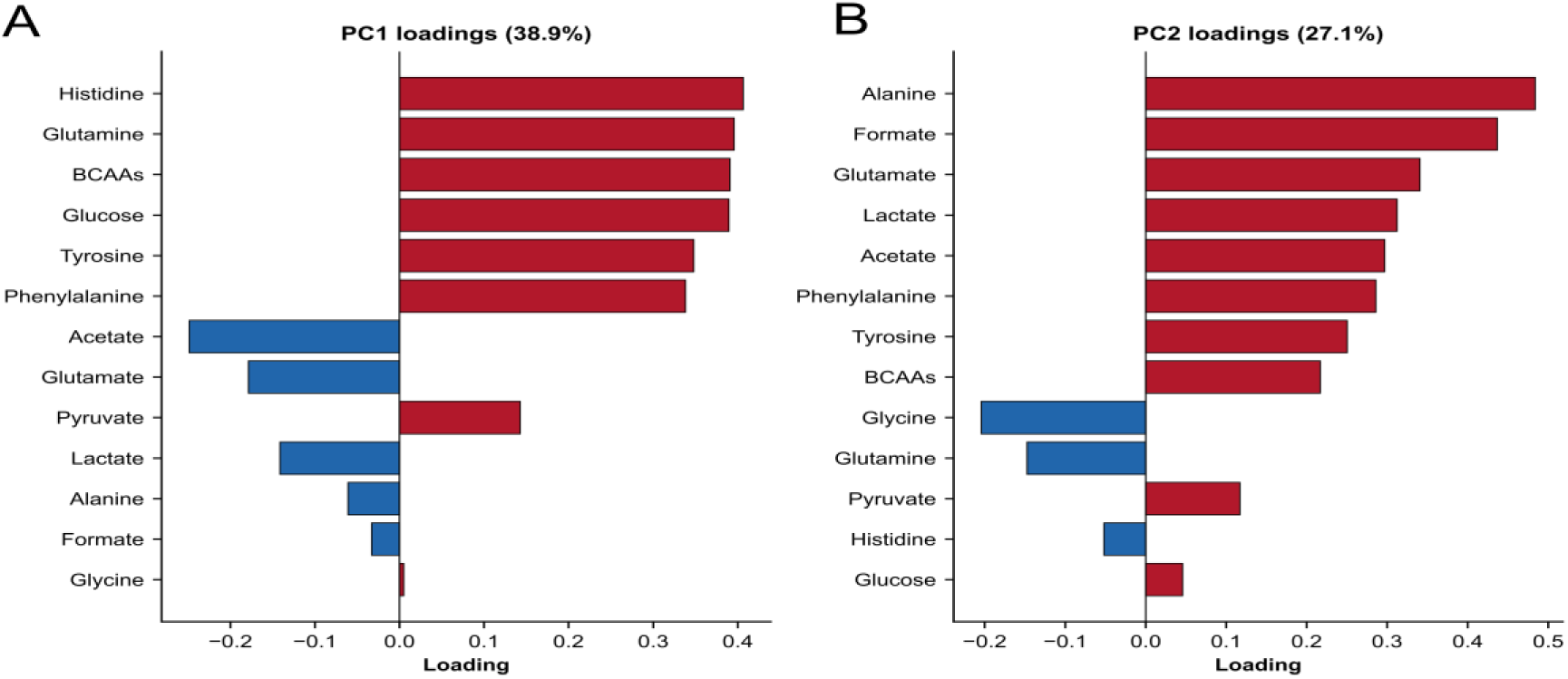
PCA loadings for PC1 and PC2. Bar charts showing the loading coefficients of the 13 quantified metabolites for (A) PC1 (38.9% of total variance) and (B) PC2 (27.1% of total variance). Red bars indicate positive loadings and blue bars indicate negative loadings. Metabolites are ordered by loading magnitude within each panel. The loading plots identify the metabolites contributing most strongly to the separation observed in the PCA score plot (Fig. 6A). Metabolic flux data were z-score normalized prior to PCA.

**Table S1.** NMR integration peaks.

| Metabolite | Integration Range (ppm) | N<sup>a</sup> |
| --- | --- | --- |
| α-D-Glucose | 5.24–5.28 | 1 |
| β-D-Glucose | 4.63–4.69 | 1 |
| Lactate | 4.10–4.18 | 1 |
| Pyruvate | 2.38–2.40 | 3 |
| Acetate | 1.937–1.946 | 3 |
| Alanine | 1.48–1.52 | 3 |
| Glycine | 3.53–3.55 | 2 |
| Glutamine | 2.10–2.18 | 2 |
| Glutamate | 2.04–2.10 | 2 |
| BCAAs (Leu+Ile+Val) | 0.85–1.07 | 18 |
| Tyrosine | 6.80–6.95 | 2 |
| Phenylalanine | 7.33–7.47 | 5 |
| Histidine | 7.77–7.80 | 1 |
| Formate | 8.45–8.50 | 1 |

<sup>a</sup> **Number of protons contributing to the integrated signal.**

